# CODEX: COunterfactual Deep learning for the *in-silico* EXploration of cancer cell line perturbations

**DOI:** 10.1101/2024.01.24.577020

**Authors:** Stefan Schrod, Tim Beißbarth, Helena U. Zacharias, Anne-Christin Hauschild, Michael Altenbuchinger

## Abstract

**Motivation:** High-throughput screens (HTS) provide a powerful tool to decipher the causal effects of chemical and genetic perturbations on cancer cell lines. Their ability to evaluate a wide spectrum of interventions, from single drugs to intricate drug combinations and CRISPR-interference, has established them as an invaluable resource for the development of novel therapeutic approaches. Nevertheless, the combinatorial complexity of potential interventions makes a comprehensive exploration intractable. Hence, prioritizing interventions for further experimental investigation becomes of utmost importance.

**Results:** We propose CODEX as a general framework for the causal modeling of HTS data, linking perturbations to their downstream consequences. CODEX relies on a stringent causal modeling strategy based on counterfactual reasoning. As such, CODEX predicts drug-specific cellular responses, comprising cell survival and molecular alterations, and facilitates the *in-silico* exploration of drug combinations. This is achieved for both bulk and single-cell HTS. We further show that CODEX provides a rationale to explore complex genetic modifications from CRISPR-interference *in silico* in single cells.

**Availability and Implementation:** Our implementation of CODEX is publicly available at https://github.com/sschrod/CODEX. All data used in this article are publicly available.

## 1 Introduction

Large-scale perturbation experiments in human cancer cell lines offer a powerful approach to connect genetic or chemical interventions with downstream effects. Such high-throughput screens (HTS) aid the identification of new drug compounds and more effective cancer treatments, and provide a way to study genomic susceptibilities in cancer [Ling et al., 2018].

This motivated several data collections and technical advancements. The Genomics of Drug Sensitivity in Cancer (GDSC) database [Iorio et al., 2016] provides response measurements of 1,001 cancer cell lines to 265 anti-cancer drugs. In comparison to individual drugs, drug combinations can have increased efficacy, and reduced toxicity and adverse side effects as a consequence of reduced dosages [Csermely et al., 2013, Al-Lazikani et al., 2012]. This motivated drug combination databases such as DrugComb [Zagidullin et al., 2019], gathering data of more than 700,000 drug combinations for more than 8000 unique compounds. Investigated downstream effects are not limited to measures of drug response. The Connectivity Map [Subramanian et al., 2017] offers more than 3,000,000 perturbed gene expression profiles using the L1000 technology (measuring the expression of 978 genes) (https://clue.io/), containing diverse genetic (shRNA, CRISPR, and overexpression), chemical and physiological perturbations. Advances in barcoding strategies enabled bulk RNA sequencing at comparatively low costs [Bush et al., 2017, Ye et al., 2018] and allowed the generation of perturbation profiles without restricting the gene space (see, e.g., PANACEA [Douglass et al., 2022]). A further essential development are high-throughput single-cell perturbation screens, providing measurements of perturbed transcriptomes of individual cells for diverse interventions. For instance, Sci-Plex was used to screen cancer cell lines exposed to different compounds and dosages at single-cell resolution [Srivatsan et al., 2020]. Genetic perturbations on a single-cell level were approached by Perturb-Seq, which uses barcoding techniques and CRISPR interference to perform genome-scale perturbation screens covering more than 1000 gene knockouts on RPE-1 and K562 cells [Replogle et al., 2022]. Further, Norman et al. [2019] explored the effects of 131 two-gene knockouts on K562 cells. Even though more than 100,000 single cells are recorded, only a small fraction of the combinatorial space could be experimentally covered, highlighting the need for computational approaches to infer novel drug combination effects and guide experimental studies.

The outlined resources differ with respect to the employed techniques, and the investigated downstream effects and interventions. However, different data types typically require tailored solutions. For instance, drug sensitivity predictions in cell lines were addressed in an NCI-DREAM challenge [Costello et al., 2014] and identified a Bayesian multitask multiple kernel learning approach to perform best. Subsequent methods built on this idea and additionally addressed the model adaptation to real tumor specimens to provide treatment response predictions in individual patients [Sharifi-Noghabi et al., 2021, He et al., 2022]. The prediction of drug synergies was pioneered by DeepSynergy [Preuer et al., 2018]. DeepSynergy predicts drug synergisms from cell-line transcriptomic data in combination with features representing compound structures. Drug response predictions in single cells were addressed by the Compositional Perturbation Autoencoder (CPA) [Lotfollahi et al., 2023], which also addressed drug combinations, by perturbing the latent representation of a Variational Autoencoder (VAE). The prediction of unseen genetic interventions and combinations thereof was addressed by GEARS [Roohani et al., 2023]. GEARS utilizes Graph Neural Networks (GNNs) to incorporate prior knowledge of gene-gene relationships into the model architecture. The generative approach, however, limits applications to a specific cell type. In summary, all outlined approaches are tailored to the problem at hand, even though they describe the common problem of predicting causal consequences of a specified intervention.

The inference of many causal actions necessitates counterfactual reasoning. For instance, in a typical clinical trial, patient outcome is recorded for a treated and a control group. By comparing the outcomes of both groups, the average treatment effect can be derived. However, a treatment that is beneficial on average may not be helpful for every individual patient. The computational challenge in predicting individual patients’ treatment responses arises from the fact that patients’ outcome can only be observed for the specific treatment they received, not for the alternative treatments they did not receive (here, the control) [Rubin, 1974]. Consequently, algorithms cannot directly learn rules for selecting the “better” treatment for each patient. Instead, they must infer the alternative, or “counterfactual”, outcomes in order to adequately assess the treatment’s relative benefit. In HTS, alternative interventions can be observed, assuming that cultures of the same cell line can be considered as technical replicates. Nevertheless, it is impractical to test every possible intervention, especially for combinations of interventions; the combinatorial complexity makes a comprehensive exploration intractable. Thus, *in-silico* approaches become necessary to prioritize interventions for further experimental investigations. To address this task, we will build on counterfactual machine learning approaches to extrapolate the space of interventions to the yet unseen “counterfactual” perturbations and cell lines.

Counterfactual deep-learning (DL) approaches turned out to be particularly promising due to their enormous flexibility. The typical strategy is to construct networks which facilitate joint representation learning across all investigated interventions, and to account for treatment specific effects via dedicated network branches [Johansson et al., 2016, Shalit et al., 2017, Yao et al., 2018, Schrod et al., 2023]. In the context of HTS data, molecular information is first aggregated in a treatment-agnostic manner to encode features of unperturbed control cells and then separated into treatment-specific representations to capture the treatment underlying molecular mechanisms. Note that a dedicated representation has to be trained for each intervention or intervention combination, and as such, downstream effects of novel combinations cannot be inferred. Alternative approaches that only consider the intervention as a predictor variable typically capture only average treatment effects and may overlook individual treatment effects. This can be explicitly seen from the comparison of an ordinary regression model, which uses a treatment variable, and a T-learner, which consists of a set of individual treatment-specific models [Shalit et al., 2017, Schrod et al., 2023]. Notably, most recent counterfactual DL approaches also take into account potential treatment selection biases as might be present in observational studies, for which regularization techniques for distributional balancing are used or adversarial learning techniques [Johansson et al., 2016, Shalit et al., 2017, Yao et al., 2018, Yoon et al., 2018, Schrod et al., 2022]. Confounding effects due to treatment biases, however, might be of less relevance in the context of HTS, where interventions are *a priori* unbiased.

Here, we will introduce CODEX (COunterfactual Deep learning for the *in-silico* EXploration of cancer cell line perturbations), which builds on counterfactual DL approaches to provide a general framework to model HTS data. In contrast to existing counterfactual DL approaches, CODEX facilitates the prediction of unseen perturbation combinations by learning from individually applied interventions and complementary combinations. CODEX can account for non-linear combinatorial effects and can incorporate prior knowledge about gene-gene relationships, such as provided by Gene Ontologies (GO). We demonstrate that CODEX can extrapolate the space of interventions to new cell lines and new treatment combinations, and via prior knowledge even to completely unseen perturbations and combinations thereof. This is illustrated for both bulk and single-cell transcriptomics data, and for the predictions of drug responses and perturbed gene-expression profiles.

## 2 Material and Methods

### 2.1 CODEX

#### 2.1.1 Model architecture

The CODEX approach (Fig. 1) is based on deep neural network architectures for counterfactual reasoning [Johansson et al., 2016, Shalit et al., 2017, Yoon et al., 2018, Schrod et al., 2022]. Let **x**_*i*_ represent a vector of unperturbed molecular features that characterizes a bulk cell line specimen *i*, or, in the context of single-cell HTS, an unperturbed single cell *i*. Throughout our analyses, gene expression levels are used as input features, but in principle, any omics data could be used. Note, however, that transcriptomics data are most readily available and have been shown to be most relevant for drug sensitivity predictions [Costello et al., 2014]. Further, let the vector 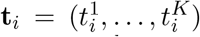 represent the perturbations applied in experiment *i*, where each element 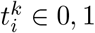 1 indicates whether intervention *k* was applied (1) or not (0). Thus, **t**_*i*_ encodes the complete set of perturbations applied in experiment *i*. In case of chemical interventions, an additional vector 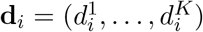 encodes drug dosages, respectively. The prediction target will be denoted as **y**_*i*_ *∈ Y*, which can be any variable or set of variables characterizing the outcome of the perturbation, such as measures of drug efficacy [Costello et al., 2014], drug synergies [Preuer et al., 2018], as well as perturbation profiles [Lotfollahi et al., 2023, Roohani et al., 2023]. Thus, CODEX requires the triplet of input information consisting of a control (the unperturbed transcriptome), the intervention (potentially associated with dosages), and the recorded downstream effects.

**Figure 1:**
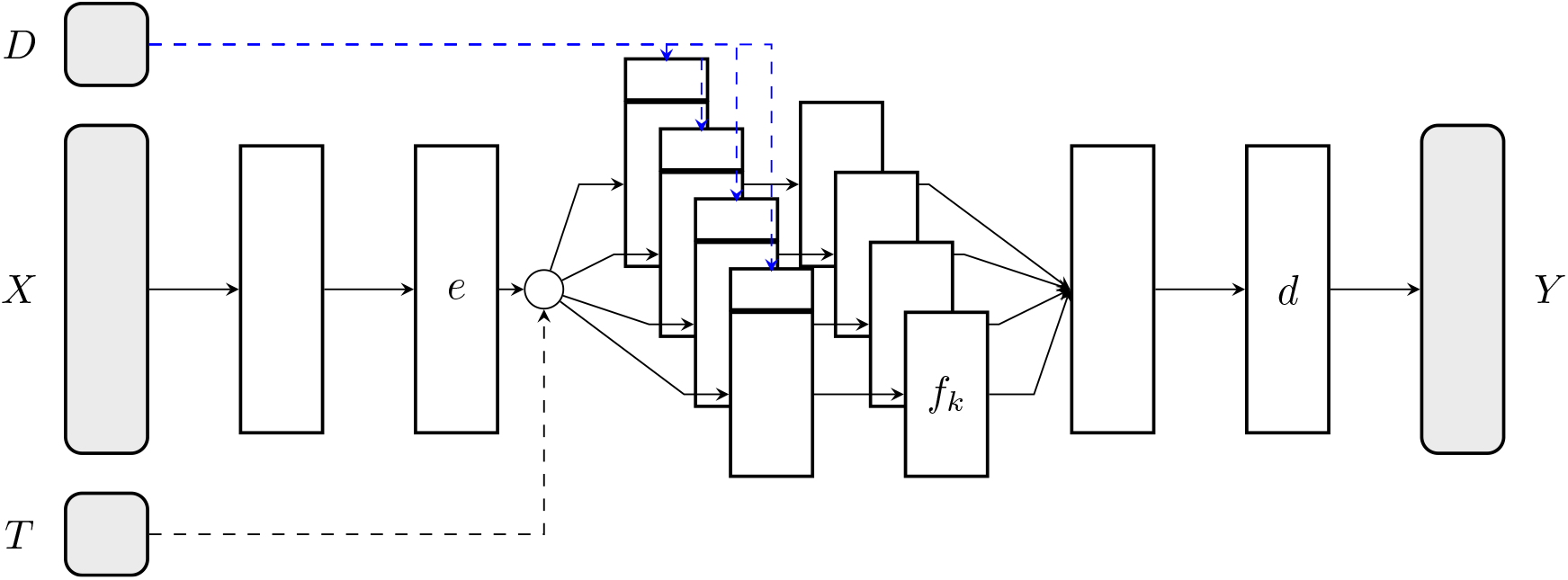
CODEX architecture used for causal representation learning: an unperturbed profile *X* is transformed to map the effect of specific interventions T, which might be associated with a dosage *D*, to the perturbed outcome *Y* . The unperturbed state is first encoded in a latent state *e*, which is passed through the respective treatment specific representations *f*_*t*_. Treatment combinations are naturally incorporated by simultaneously propagating profiles through respective treatment arms. Further, if dosage specific interventions are used, the dosage is incorporated as input variable to the intervention specific layers. Finally, the individual effects are aggregated and combined by a shared decoder *d* to model the perturbed outcome.

Causal inference models such as [Shalit et al., 2017] do not incorporate the action as a covariate but rather as a structural parameter in order to train intervention specific data representations. CODEX builds on this concept. Specifically, we first train a latent shared embedding of the initial state *e* : 𝕩 *→* ℝ^*d*^ to reduce the dimensionality of the problem and to construct prediction-relevant features. This embedding is then passed through intervention specific layers (a mapping *f*_*k*_ : ℝ^*d*^ *→* ℝ^*d*^) to account for the molecular downstream effects induced by intervention *k*. Importantly, we do not introduce individual representations for combinations of treatments. While this would be a direct adaptation of counterfactual DL models like those of [Shalit et al., 2017], it would significantly inflate the parameter space and hinder the inference of unseen combinations. This is because each treatment combination would require observations to train the respective network branch and to facilitate any kind of predictions, making existing causal inference models impracticable for complex HTS data. Thus, we pass the latent variables *e*(*X*) through a set of active network branches corresponding to the respective single interventions. Subsequently, they are passed through additional joint layers – a decoder *d* : ℝ^*d*^ *→* 𝕐 – to reconstruct the downstream effects. In case of multiple simultaneously applied interventions, the corresponding latent representations are aggregated before the combinatorial effect is deciphered by the decoder. One should note that, even though the effects are linearly aggregated on a latent state of the model, the decoder naturally captures non-linear combinatorial effects. In summary, the CODEX mapping reads:

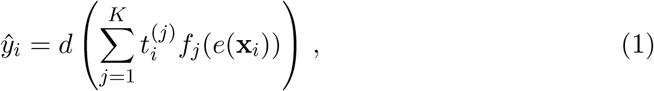

which implicitly sums all active treatment branches via the coefficients 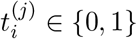.

#### 2.1.2 CODEX for drug-synergy prediction

CODEX is a general architecture to model causal effects and their combinations, and as such, enables predictions of diverse outcome measures. In this work, we focus on the prediction of drug synergies and the reconstruction of perturbed gene expression profiles. Drug synergies are commonly expressed in terms of synergy scores *Y ∈* ℝ, which quantify the difference between experimentally tested response surfaces and theoretical models combining individual treatments naively, such as Loewe Additivity [Loewe, 1953] or Bliss independence [Bliss, 1939]. In this work, we focus on the Zero Interaction Potency (ZIP) score, which takes advantage of both the Loewe Additivity and the Bliss independence model [Yadav et al., 2015]. To model drug synergies via CODEX, we map the decoder to a Mean Squared Error (MSE) loss

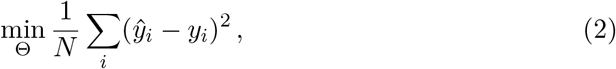

where the parameter space is given by Θ = [*θ*_*e*_, *θ*_*ft*_, *θ*_*d*_]. The applied interventions are implicit in Eq. (1). The full model architecture and the hyper-parameter search space are given in the Supplementary Materials.

#### 2.1.3 CODEX for the prediction of single-cell perturbation profiles

Synergy scores are summary statistics of potentially complex molecular downstream effects. In recent years, downstream effects have become increasingly available in terms of perturbed transcriptomes in both bulks and single cells [Subramanian et al., 2017]. We focus on the latter with *y*_*i*_ = (*y*_*i*1_, …, *y*_*ip*_) ∈ ℝ^*p*^ corresponding to a perturbed transcriptome with *y*_*ij*_ representing the expression of gene *j* in cell *i*. Let 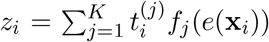 which is the linear combination of the latent variables representing active treatment branches in cell *i*. CODEX then uses a Gaussian loss which quantifies the accuracy of predictions of *y*_*i*_ together with respective gene variances,

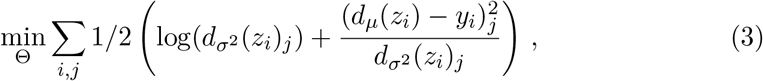

where *d*_*μ*_(*z*_*i*_)_*j*_ maps to the mean expression of gene *j* and 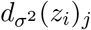 infers the variance of the estimates, given the respective treatment or treatment combination. We further imposed that 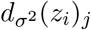 is larger than *ϵ* = 10^*−*6^ to increase numerical stability.

#### 2.1.4 CODEX for the prediction of drug dosage effects

Different drug dosages add to the combinatorial complexity of intervention experiments and can introduce complex non-linear dependencies. In principle, dosage effects can be incorporated into CODEX via different treatment branches. This, however, would prohibit the extrapolation to unseen drug dosages and would substantially increase model complexity. Therefore, rather than introducing additional dose representations, CODEX incorporates dosage information as an additional categorical feature 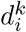 which adds to the first layer of each treatment specific branch *k* (Fig. 1).

#### 2.1.5 CODEX for the prediction of unobserved perturbations

CODEX infers a distinct model branch for each individual perturbation, effectively reducing the number of model representations to the number of distinct perturbations. However, it does not require that those perturbations were independently observed. Rather, the training data can comprise perturbations observed in combinations. Consequently, each combination adds information to decipher the individual model branches. We make use of this feature and introduce a weighting scheme to share information among perturbations.

We illustrate this concept for CRISPR interference screens, where the individual perturbations correspond to silenced genes. In this case, we use gene similarities derived by [Roohani et al., 2023], which aggregate information from GO [Consortium, 2004]. The basic idea is to compute the Jaccard index between a pair of genes *j,j*^*′*^ as 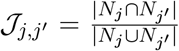 where *N*_*u*_ is the set of pathways containing gene *u*. This is the fraction of shared pathways between two genes.

Consider an unobserved gene knockout of gene *j*^*′*^. Then, we can construct the treatment proxy model by setting 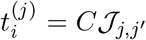 where *C* is determined by normalization 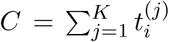.In the case of single unobserved perturbations, the proxy vector is used as is, otherwise, respective proxy vectors and observed treatment vectors are added up. For instance, let (1, 0, 0, 0, 0) be a treatment vector, where the observed interventions are indicated by the 1 at the first position. Further, let (0, 0.4, 0.35, 0.25, 0) be a normalized proxy treatment vector, determined as outlined. Then, we evaluated CODEX with the sum of both, **t**_*i*_ = (1, 0.4, 0.35, 0.25, 0). Thus, in summary, CODEX passes the data through treatment branches which likely resemble the unobserved perturbation.

For more details about the implementation and the used network architecture refer to the Supplementary Materials.

### 2.2 Competing methods

#### 2.2.1 Algorithms for drug-synergy predictions in cell lines

We compared CODEX to

- TreeCombo [Janizek et al., 2018], which is a tree-based approach using Extreme Gradient Boosting (XGBoost),
- DeepSynergy [Preuer et al., 2018], a dense feed forward neural network to predict synergy scores,
- MatchMaker [Kuru et al., 2021], which is inspired by DeepSynergy but additionally splits the network into two parts representing the two different interventions, and
- MARSY [El Khili et al., 2023], which uses a drug-drug representation network combined with the latent representation learner of DeepSynergy.

In contrast to CODEX, the competing models additionally use chemical drug encodings as input.

#### 2.2.2 Algorithms for the prediction of post-perturbation profiles

We compared CODEX to:

- Random Baseline: in line with Lotfollahi et al. [2023], we implemented a random baseline to assess the relative benefit of CODEX in the context of dosage extrapolations.
- Linear baseline: we implemented a linear baseline which simply averages the downstream predictions of individual perturbation to predict the effect of perturbation combinations.
- Gene Regulatory Network (GRN): GRN, as implemented by Roohani et al. [2023], infers a GRN to linearly propagate the effect of gene perturbations.
- Compositional Autoencoder (CPA) [Lotfollahi et al., 2023]: CPA is based on a Variational Autoencoder (VAE) architecture trained to encode both control and perturbation profiles. It encodes different perturbations and dosages in a latent space. This space is made indistinguishable with respect to the different interventions using an adversarial discriminator. Downstream effects are predicted by (1) encoding control cells and (2) decoding them with respective interventions activated in the latent space.
- Graph-Enhanced gene Activation and Repression Simulator (GEARS) [Roohani et al., 2023]: GEARS is a generative approach assuming a single type of cell. It uses two separate Graph Neural Networks (GNNs) to encode additional prior knowledge about gene-gene relationships and perturbation relationships. The first GNN embeds the unperturbed state using a gene co-expression knowledge graph, and the second GNN learns perturbation embeddings using a graph derived from GO. The states are combined using the respective perturbations, and a feed-forward decoder is used to reconstruct the post-perturbation gene expression.
- Linear CODEX: lin-CODEX removes the non-linear effect combination from CODEX. This baseline is included to illustrate the benefit of the effect decoder and thus serves as an ablation study. For model inference, the trained CODEX model is evaluated only for individual model branches and subsequently averaged to receive predictions for respective treatment combinations.

## 3 Results

### 3.1 CODEX improves drug-synergy predictions in cancer cell lines

DrugComb [Zagidullin et al., 2019] provides more than 700.000 recorded drug combinations for 8379 different drugs on 2320 different tissues, making it an invaluable resource to explore drug combinations. However, the combinatorial complexity prohibits a comprehensive experimental exploration and asks for computational solutions. For instance, exploring all combinations of two drugs in all tissues would correspond to *∼* 10^11^ experiments. Following El Khili et al. [2023], we extracted 670 unique drugs and a set of 2353 corresponding drug pairs on 75 selected cancer cell lines. Synergy effects were extracted from DrugComb [Zagidullin et al., 2019], and the normalized untreated cancer cell lines were retrieved from CellMiner [Reinhold et al., 2012]. Lowly expressed genes with log_2_(RPKM+1) *<* 1 and a variance smaller than 0.8 were excluded, resulting in a final set of 4639 features. For validation, we followed El Khili et al. [2023], and performed two different settings:

1. a 5-fold cross-validation, where randomly triplets of cell line-drug-drug were selected for testing,
2. a stratified 5-fold cross-validation strategy, where individual treatment combinations were removed from the training, meaning that the testing fold contains unseen drug combinations.

We evaluated the Zero Interaction Potency (ZIP), which incorporates Loewe Additivity [Loewe, 1953] and the Bliss independence [Bliss, 1939] score. All baselines were extracted from [El Khili et al., 2023].

### Performance comparison

The results are given in Table 1. We evaluated three performance measures: the Spearman Correlation Coefficient (SCC), the Pearson Correlation Coefficient (PCC) and the MSE between ground truth and predictions. In the first scenario, the test set contained drug-drug combinations which were already part of the training data, although they were not seen in the same cell line. CODEX yielded the highest PCC and lowest MSEs between predictions and ground truth values and was only out-competed by MARSY with respect to SCC. The remaining competitors performed substantially worse than both approaches.

**Table 1:** Mean 5-fold cross-validation performance for the prediction of ZIP synergy scores, in terms of Spearman Correlation Coefficient (SCC), Pearson Correlation Coefficient (PCC) and Mean Squared Error (MSE) for unseen cell lines (1) and unseen drug combinations (2).

| Model | Setting (1) |  |  | Setting (2) |  |  |
| --- | --- | --- | --- | --- | --- | --- |
|  | SCC | PCC | RMSE | SCC | PCC | RMSE |
| CODEX | 0.752 ($\pm 0.004$) | 0.889 ($\pm 0.005$) | 5.31 ($\pm 0.18$) | 0.725 ($\pm 0.005$) | 0.877 ($\pm 0.005$) | 5.58 ($\pm 0.17$) |
| MARSY | 0.780 ($\pm 0.010$) | 0.886 ($\pm 0.005$) | 5.36 ($\pm 0.19$) | 0.749 ($\pm 0.011$) | 0.875 ($\pm 0.005$) | 5.62 ($\pm 0.15$) |
| MatchMaker | 0.742 ($\pm 0.004$) | 0.873 ($\pm 0.006$) | 6.11 ($\pm 0.32$) | 0.720 ($\pm 0.006$) | 0.864 ($\pm 0.007$) | 6.23 ($\pm 0.12$) |
| TreeCombo | 0.737 ($\pm 0.004$) | 0.870 ($\pm 0.003$) | 5.73 ($\pm 0.17$) | 0.689 ($\pm 0.005$) | 0.856 ($\pm 0.006$) | 6.00 ($\pm 0.15$) |
| DeepSynergy | 0.701 ($\pm 0.003$) | 0.869 ($\pm 0.004$) | 5.78 ($\pm 0.18$) | 0.676 ($\pm 0.006$) | 0.860 ($\pm 0.008$) | 5.95 ($\pm 0.16$) |

Next, we evaluated the more challenging scenario of validation experiment (2), where the test fold contains only unseen drug combinations. As expected, all approaches performed worse than in the validation experiment (1). However, both CODEX and MARSY still achieve reasonable performance with PCCs *>* 0.87. As previously, CODEX performed best with respect to PCC and MSE and MARSY with respect to SCC. The same holds true considering an alternative synergy score *S*_mean_, where MARSY slightly out-competed CODEX, while the other approaches showed inferior performance (refer to Supplementary Materials Table 1).

### 3.2 CODEX improves dose-response predictions in single-cell data

We studied CODEX’s ability to predict dose specific responses in single-cell perturbation data. We used the Sciplex2 data [Srivatsan et al., 2020], which contain 12656 post-perturbation transcriptomic profiles of A549 human lung adenocarcinoma cells measured for 4 different perturbations (Vorinostat, BMS-34554, Dexamethasone and Nutlin3a) and 7 different concentrations (in total 28 drug-dose combinations). We used the preprocessed data provided by Lotfollahi et al. [2023], which were normalized and log transformed and limited to the 5000 most variable genes. Accordingly, we left out the second highest treatment concentration and tested CODEX ability to reproduce the dose response curves and to interpolate to unseen concentrations.

### Performance comparison

Fig. 2 shows dose-response curves (response versus dosage) for the top three Differentially Expressed Genes (DEGs) for each treatment. Ground truth values (solid lines and error bars), with dots representing individual measurements, are contrasted with predictions by CPA (dotted lines) and CODEX (dashed lines). The second highest dosage was hold out for testing and is highlighted by a vertical dashed line. CODEX achieves a substantially better description of both training and test dosages, capturing also non-linear dependencies (Fig. 2). This is also supported by considering the reconstruction performance in terms of the coefficient of determination *R*^2^ (4), for all genes (blue) and for the top 50 DEGs (orange). CODEX performed best for Dexamethasone, Nutlin-3a, Vorinostat and the linear baseline for BMS-34554, where substantial improvements on the top 50 DEGs were observed for the drugs Vorinostat (0.94 vs. 0.85 for CODEX vs. CPA), and Nutlin-3a (0.84 vs. 0.81 for CODEX vs. CPA). On average, we observed a median performance gain of 9.1% for the top 50 DEGs compared to CPA (Supplementary Fig. 1A). Furthermore, considering all dosages (both validation and ood data), CODEX showed significantly improved *R*^2^ values compared to CPA, and a median performance increase of 11.5% for the top 50 DEGs (Supplementary Fig. 1B).

**Figure 2:**
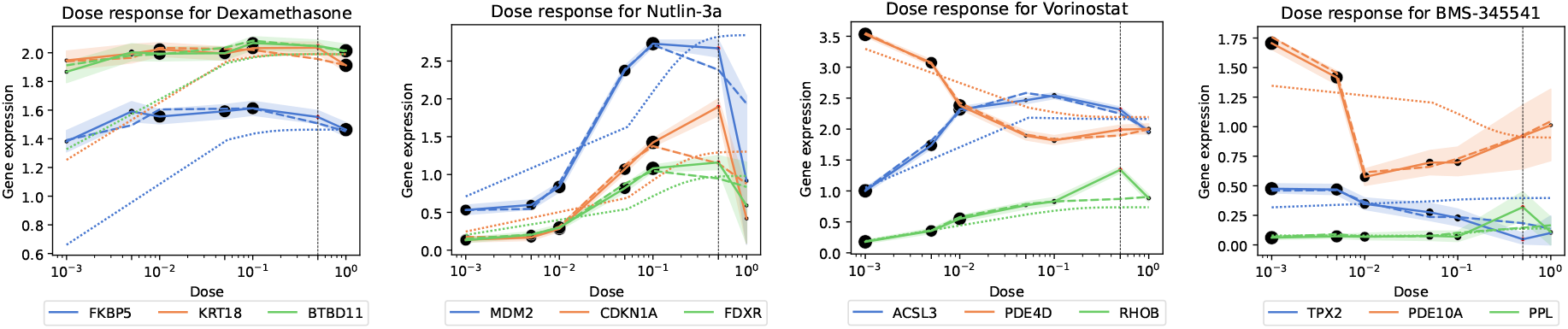
Dose-dependent reconstruction of the gene expression levels of the top 3 DEGs for each of the treatments. Ground truth values (solid lines and error bars), with the size of the dots representing the number of available samples for each measurement, are contrasted with predictions by CPA (dotted lines) and CODEX (dashed lines). The dashed vertical line indicates the dosage left out for testing.

**Figure 3:**
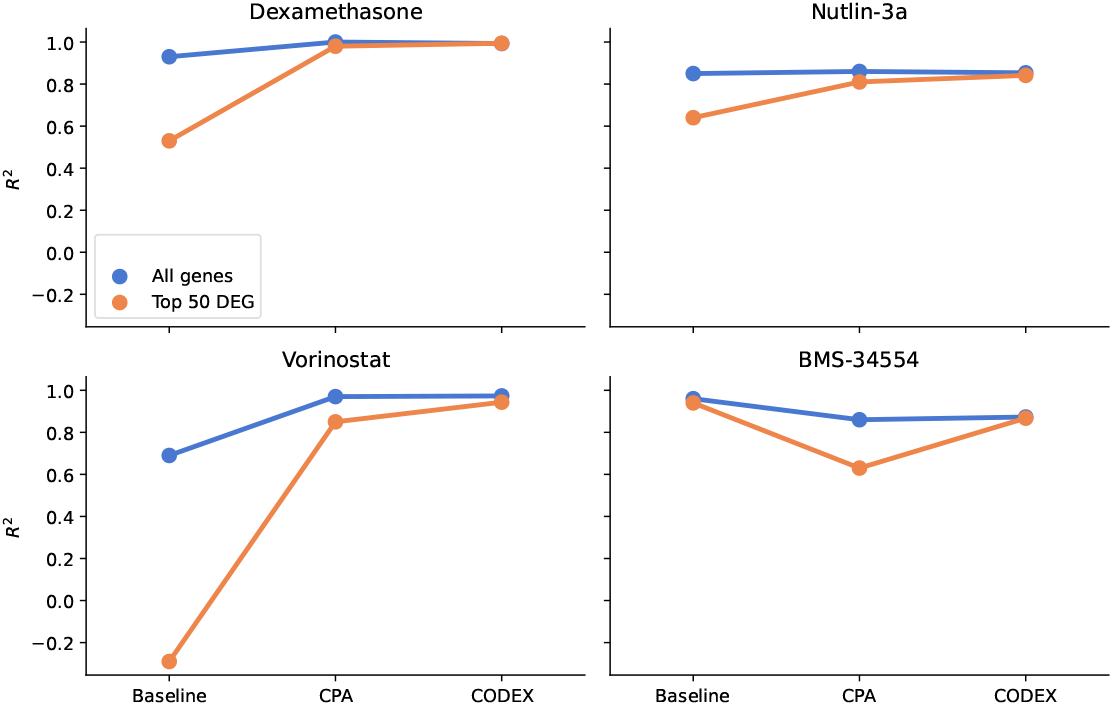
*R*^2^ reconstruction performance of the mean post perturbation gene expression of all genes (blue) and the top 50 DEGs (orange) obtained for the second highest dose (left out for testing) on Sciplex2.

### 3.3 CODEX can predict molecular downstream effects of drug combinations in single cells

We further investigated molecular downstream effects in terms of perturbed transcriptomes of 13 anticancer drugs on A549 cells obtained from the combinatorial sci-Plex (Combosciplex) assay [Lotfollahi et al., 2023]. The data comprise 18 different anti-cancer medications evaluated on 63.430 single cells, including a total of 25 unique drug combinations and 7 individually observed drug perturbations (for an overview, see Supplementary Table 5). Similar to the Sciplex2 data, Combosciplex was normalized, log-transformed, and restricted to the 5000 most variable genes. To study the ability of CODEX to infer the effect of unseen drug combinations, we excluded four combinations during model development and evaluated the reconstruction of perturbed transcriptomes using *R*^2^ scores. We compared CODEX to the linear baseline, which averages the observed single effects, CPA [Lotfollahi et al., 2023], and lin-CODEX.

### Performance comparison

Considering median *R*^2^ values, we observed that CODEX performs best with an increase of 5.5% on the top 50 DEGs compared to CPA (Supplementary Fig. 2). The performance resolved for the individual hold-out treatment combinations is shown in Fig. 4. There, already lin-CODEX is able to improve predictions compared to the linear baseline. However, it is outperformed by both CPA and CODEX, suggesting a crucial role for non-linear decodings. The performance gains of CODEX were mainly attributed to the combinations including Alvespimicin (Fig. 4). Those were weakly supported by the training data with significantly different effects than observed during training [Lotfollahi et al., 2023]. This is further illustrated in a UMAP representation, where the left-out treatment combinations containing Alvespimin (purple triangles, with blue circles) are separated from the majority of the training combinations (red circles), both on the latent space and the final predictions (Fig. 5A and B).

**Figure 4:**
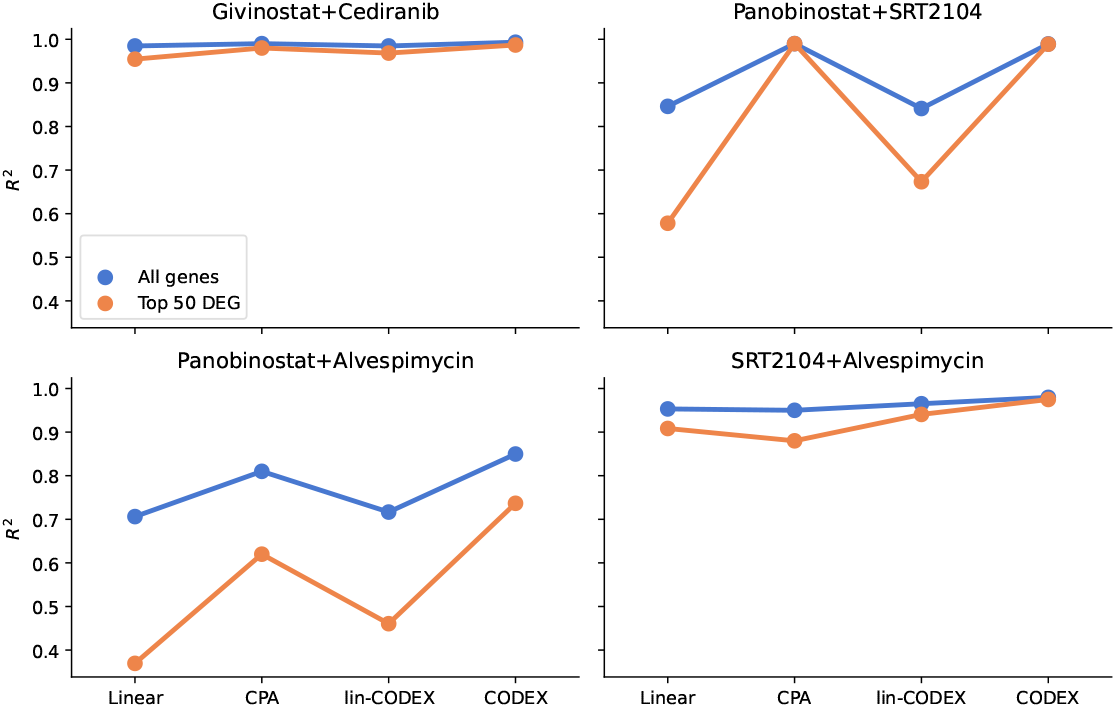
*R*^2^ reconstruction performance of the mean post perturbation gene expression profiles for all genes (blue) and top 50 DEGs (orange) obtained for the four held out treatment combinations on the Combosciplex data.

**Figure 5:**
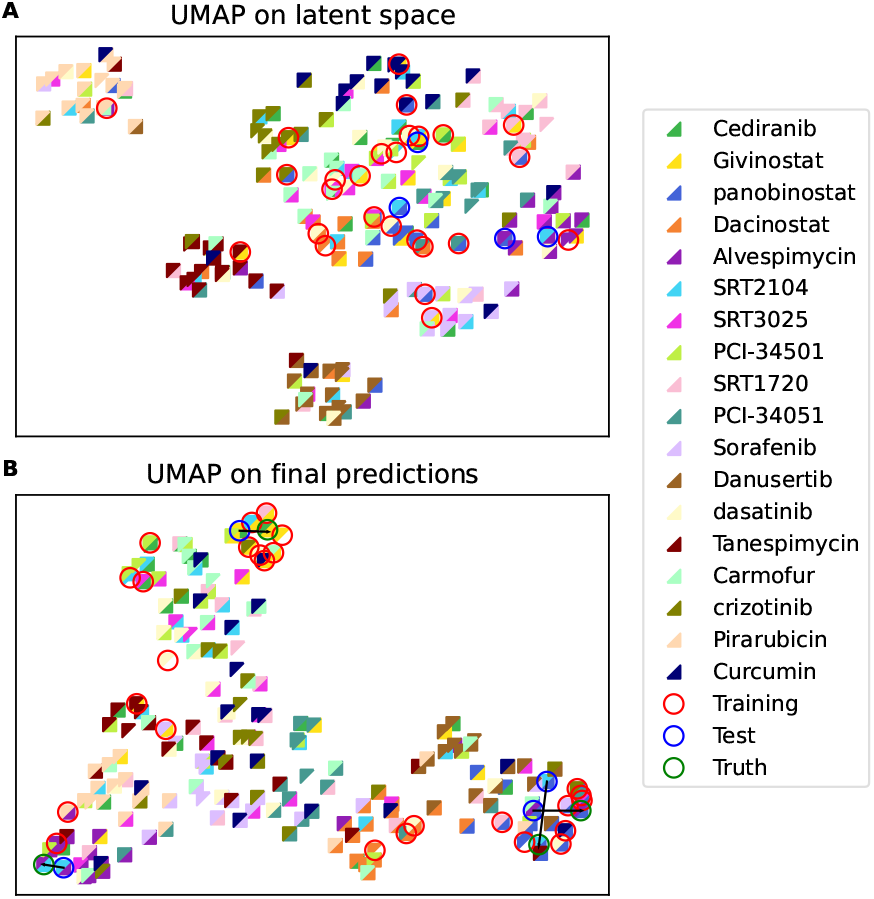
UMAP of the combined latent representation *z* of CODEX (A) and of its final predictions (B) for all possible treatment combinations of the Combosciplex data. Each colored triangle represents a treatment component, and the full squares represent their respective combination. Training combinations are circled in red, test combinations in blue, and the ground truth for the unobserved combinations in green (only for final predictions).

The Combosciplex data comprise only a subset of all possible drug combinations. We next used CODEX to infer all remaining unobserved drug combinations and visualized the results using a UMAP representation on the latent linear effects (*z*_*i*_) and on the final predictions (top and bottom of Fig. 5, respectively). UMAP on the latent representation reveals distinct clusters associated with the dominant effects attributed to Pirarubicin, Danusertib, Tanespimycin, and Sorafenib. Those are not revealed on the final predictions (bottom Fig. 5), suggesting that the non-linear adjustments of the decoder regulate the treatment effect sizes. We further confirmed that out-of-sample predictions (blue circles) are in near vicinity of respective ground truths (green circles), with one-to-one correspondence indicated by connecting black lines.

### 3.4 CODEX facilitates predictions of combined gene knock-out perturbations from CRISPRi in single cells

Finally, we used CODEX to explore genetic perturbations implemented via CRISPRi. CRISPRi facilitates the targeted silencing of genes and has become increasingly feasible in large-scale single-cell perturbation screens in recent years. For instance, Norman et al. [2019] proposed Perturb-seq to perform single-cell pooled CRISPRi screens and provided data containing a total of 284 unique knock-out conditions, comprising 131 unique two-gene knockouts, on 108.000 single-cells from K562 cancer cell line [Norman et al., 2019]. In the first experiment, we selected test combinations where the individual perturbations were part of the training data. To further guarantee a fair comparison to state-of-the-art competitors, we followed Roohani et al. [2023] and implemented the same 5 test-training splits using the same feature set of 5045 genes. We assessed the performance in reconstructing perturbed profiles in terms of normalized MSE on the top 20 DEGs (normalized using the reconstruction error of the random baseline) and PCC, where we evaluated the reconstructed effect on all genes (not considering the control background) and the reconstruction of the top 20 DEGs.

### Performance comparisons

Also, considering this application, CODEX substantially improves the recently suggested state-of-the-art solutions CPA and GEARS and by far out-competes GRN. Considering PCC for the top 20 DEGs (Fig. 6C), we observed a median value of 0.98 for CODEX compared to 0.96 for lin-CODEX, 0.91 for CPA, 0.92 for GEARS, and 0.82 for GRN. This is consistent with the performance in terms of normalized MSEs (Fig. 6A) and by considering PCC for all genes (Fig. 6B). To further substantiate these findings, we performed an additional validation experiment corresponding to the setting of [Lotfollahi et al., 2023], where each combination of perturbations is left out in one of 13 splits, and evaluated the reconstruction performance in terms of the *R*^2^ score on all genes and on the top 50 DEGs, in line with [Lotfollahi et al., 2023]. These results demonstrate that in this comparison, CODEX out-competes the other methods significantly (Supplementary Fig. 4).

**Figure 6:**
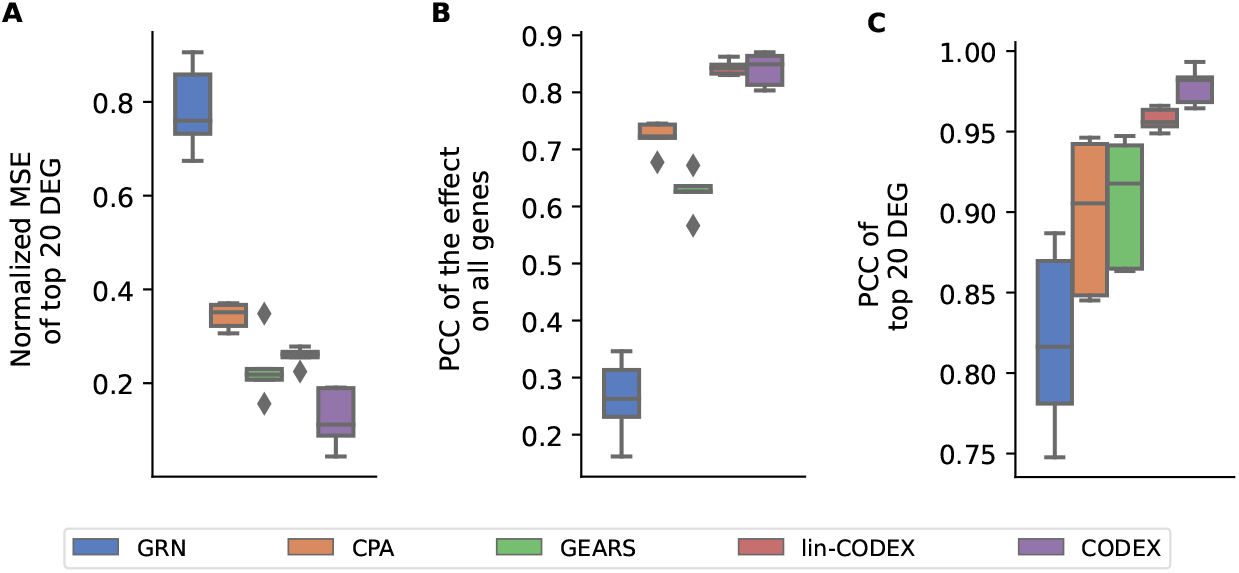
Reconstruction performance of unseen perturbation combinations on the Norman et al. [2019] data.

### 3.5 CODEX facilitates predictions of unobserved gene knock-outs from CRISPRi in single cells

CODEX can predict downstream effects of unseen perturbations, combinations thereof with observed perturbations, as well as combinations of unseen perturbations only. This is achieved by proxy models resembling unseen perturbations. These proxies were established by a weighting scheme summarizing related gene perturbations (see Methods). To assess CODEX’s capability to predict unseen perturbation, we performed three additional experiments using the Norman et al. [2019] dataset, where we inferred the effect of left-out perturbations for a single missing perturbation (0/1 seen), a missing perturbation in a pair (1/2 seen), and two missing perturbations in a pair (0/2 seen). Due to the limited number of single gene perturbations (105), we tested CODEX on two additional genetic perturbation screens generated by Replogle et al. [2022], comprising a total of 1,543 RPE-1 and 1,092 K562 single genetic perturbations, with 175, 398 and 192, 648 measured single cells, respectively. We again adapted the experimental setup of Roohani et al. [2023] and compared the reconstruction error based on 5 identical sets of held-out perturbations. Performance was again assessed using normalized MSE on the top 20 DEGs, PCC of the effect on all genes, and PCC on the top 20 genes.

### Performance comparison

For the Norman et al. [2019] data, we observed that GEARS yielded the lowest median MSE in all three settings (Fig. 7A). However, CODEX consistently out-competes all other methods with respect to PCC (Fig. 7B and C). There, CODEX improves the median PCC on the top 20 DEGs from 0.90 to 0.92 for 0/1 seen, 0.83 to 0.93 for 1/2 seen, and 0.79 to 0.89 for 0/2 seen compared to the second best performing model (GEARS).

**Figure 7:**
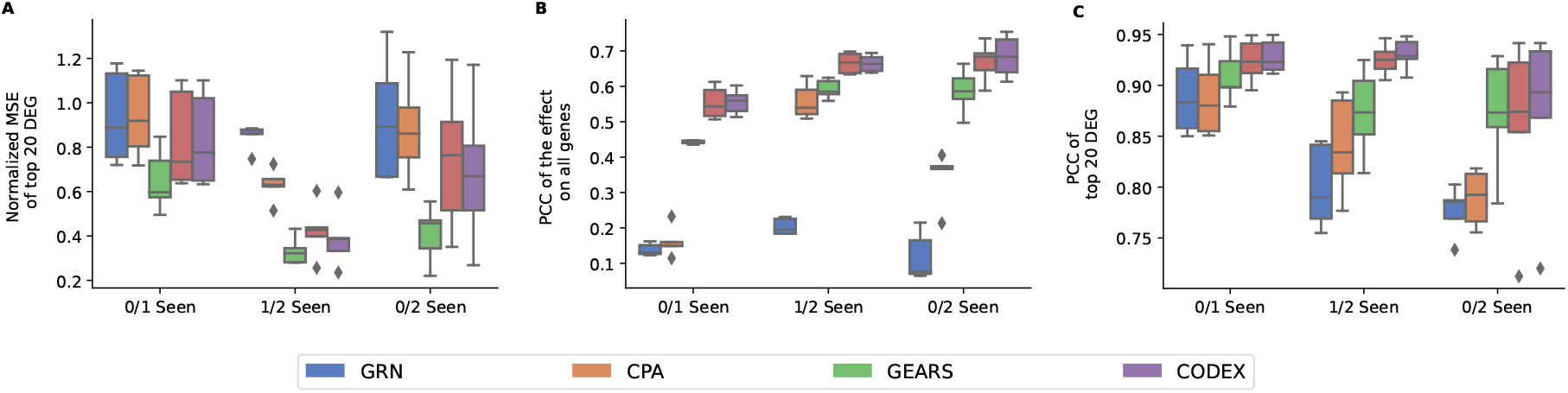
Reconstruction performance of unseen perturbations for varying degrees of difficulty, 0/1 seen, 1/2 seen, and 0/2 seen on the Norman et al. [2019] data.

On the Replogle et al. [2022] data, CODEX performs best on all compared measures for both K562 and RPE-1 cells (Fig. 8a-c). On K569 cells, CODEX improves the median PCC from 0.31 to 0.49, and on RPE-1 cells from 0.53 to 0.61, compared to GEARS (Fig. 8b). In comparison, the performance of CPA and GRN is highly compromised with respect to MSE (Fig. 8a) and PCC on all genes (Fig. 8b). These findings suggest that when many single perturbations are known, gene graph proxies can be highly efficient. This holds true for both CODEX and GEARS. For CODEX, however, this aspect seems to be even more beneficial.

**Figure 8:**
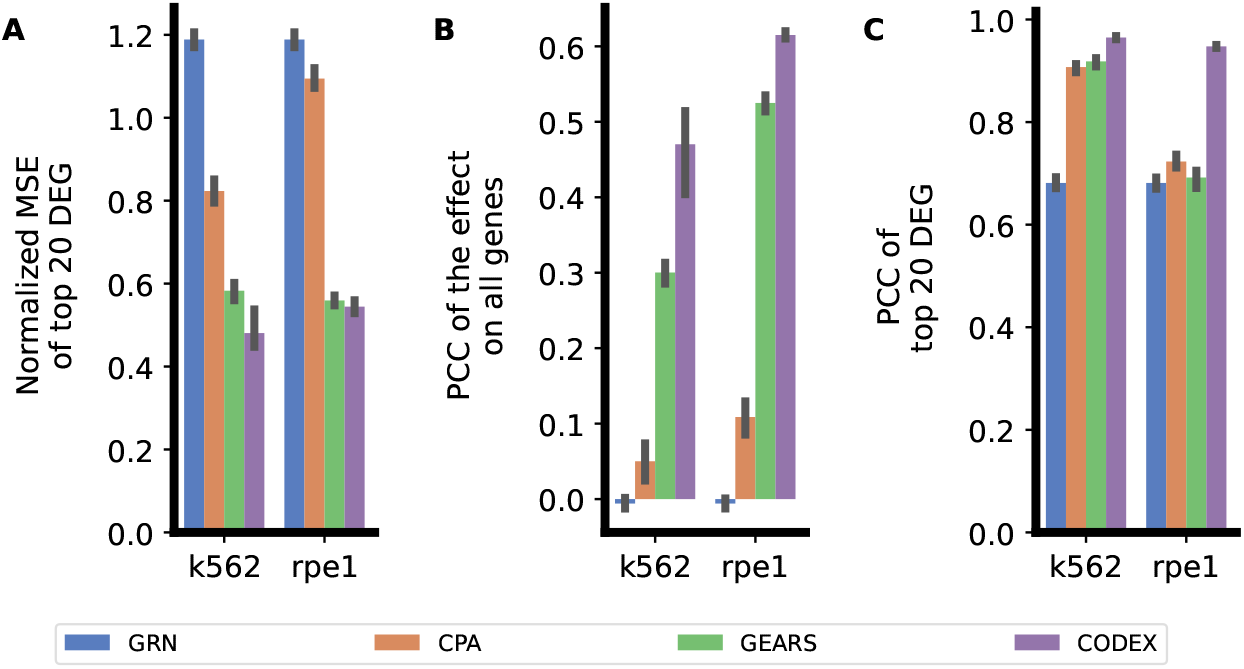
Reconstruction performance for an unseen single perturbation on single cells from K562 and RPE-1 cell lines [Replogle et al., 2022].

## 4 Conclusion

We proposed CODEX as a general framework to model high-throughput perturbation experiments. CODEX naturally facilitates diverse causal prediction tasks, can learn non-linear effect combinations, and can model different intervention types. This was shown for both chemical and genetic perturbation combinations. Moreover, we suggested a weighting scheme to perform predictions for completely unseen perturbations. Thus, applications of CODEX are rich and the outlined performance comparisons suggest that CODEX offers highly competitive performance across diverse applications.

CODEX builds on counterfactual reasoning and implements different perturbations via distinct model representations, while most state-of-the-art approaches include perturbations as distinct variables Lotfollahi et al. [2023], Roohani et al. [2023] or represent them using chemical embeddings [Preuer et al., 2018, Kuru et al., 2021, El Khili et al., 2023]. Causal modeling of representations increases model complexity, but has the benefit that highly complex downstream effects can be captured through the model architecture. This might be particularly relevant for the prediction of complex phenotypes, as our empirical results for the prediction of perturbed single-cell transcriptomes suggest. To enable out-of-distribution predictions for unseen drug combinations, different perturbation branches are combined. This has two advantages. First, subsequent network layers can account for potential non-linearities. Those were repeatedly shown to be key to the outlined prediction tasks as supported by our comparison to the linear CODEX baseline (lin-CODEX). Second, and most importantly, CODEX can learn the individual perturbation from combinations of perturbations. Given the combinatorial complexity of high-throughput perturbation screens, it is not tractable to comprehensively explore the space of perturbations and their downstream effects experimentally. However, CODEX makes use of redundancy, and might further improve extrapolation performance as perturbations are captured in more and more diverse combinations, not even limited to pairs of perturbations. This will be crucial to maximize the use of upcoming, potentially more comprehensive perturbation screens.

CODEX has a number of limitations. First, the main issue that afflicts all prediction algorithms of such a kind is that *in vitro* cell line experiments provide only a very limited picture of the mechanisms taking place *in vivo*. It will be key to further consider the *in vivo* model transfer. Multiple approaches were already suggested, comprising Velodrome [Sharifi-Noghabi et al., 2021] and CODE-AE [He et al., 2022]. However, they mainly rely on a measure of distributional similarity and do not address that biological processes between human and cell lines differ, with the former comprising, e.g., complex cellular interactions taking place in the tumor microenvironment. Second, nowadays perturbation experiments are destructive, meaning that we never observe an individual cell before and after the intervention. Observing the latter might substantially deepen our understanding of molecular downstream effects. In this direction, e.g., Bunne et al. [2023] and Dong et al. [2023], attempt to identify counterfactual cell pairs using optimal transport. However, a combination of those methods with CODEX requires future research, addressing, e.g., potential biases from cell-pair selection.

In summary, CODEX provides a highly potent framework to extrapolate the space of interventions in high throughput perturbation experiments to unseen interventions and combinations thereof. Empirical results suggest that it substantially enhances out-of-distribution predictions and applies to diverse prediction tasks, suggesting rich applications in pharmacogenomics.

## Data and availability

All data used in this article are publicly available. The Gene Expression Omnibus accession numbers to obtain the raw datasets are: Srivatsan et al. [2020] GSM4150378, Lotfollahi et al. [2023] GSE206741, Norman et al. [2019] GSE133344. The data from Replogle et al. [2022] are available from https://doi.org/10.25452/figshare.plus.20022944 and the prepossessed drug-synergy data from https://github.com/Emad-COMBINE-lab/MARSY/tree/main/data [El Khili et al., 2023]. Further information and code to reproduce the experiments are provided in the CODEX repository at https://github.com/sschrod/CODEX.

## References

Bissan Al-Lazikani, Udai Banerji, and Paul Workman. Combinatorial drug therapy for cancer in the post-genomic era. Nature biotechnology, 30(7):679–692, 2012.

Chester I Bliss. The toxicity of poisons applied jointly 1. Annals of applied biology, 26(3):585–615, 1939.

Charlotte Bunne, Stefan G Stark, Gabriele Gut, Jacobo Sarabia Del Castillo, Mitch Levesque, Kjong-Van Lehmann, Lucas Pelkmans, Andreas Krause, and Gunnar Rätsch. Learning single-cell perturbation responses using neural optimal transport. Nature methods, pages 1–10, 2023.

Erin C Bush, Forest Ray, Mariano J Alvarez, Ronald Realubit, Hai Li, Charles Karan, Andrea Califano, and Peter A Sims. Plate-seq for genome-wide regulatory network analysis of high-throughput screens. Nature communications, 8(1):105, 2017.

Gene Ontology Consortium. The gene ontology (go) database and informatics resource. Nucleic acids research, 32(suppl_1):D258–D261, 2004.

James C Costello, Laura M Heiser, Elisabeth Georgii, Mehmet Gönen, Michael P Menden, Nicholas J Wang, Mukesh Bansal, Muhammad Ammad-Ud-Din, Petteri Hintsanen, Suleiman A Khan, et al. A community effort to assess and improve drug sensitivity prediction algorithms. Nature biotechnology, 32(12):1202–1212, 2014.

Peter Csermely, Tamás Korcsmáros, Huba JM Kiss, Gábor London, and Ruth Nussinov. Structure and dynamics of molecular networks: a novel paradigm of drug discovery: a comprehensive review. Pharmacology & therapeutics, 138(3):333–408, 2013.

Mingze Dong, Bao Wang, Jessica Wei, Antonio H de O. Fonseca, Curtis J Perry, Alexander Frey, Feriel Ouerghi, Ellen F Foxman, Jeffrey J Ishizuka, Rahul M Dhodapkar, et al. Causal identification of single-cell experimental perturbation effects with cinema-ot. Nature Methods, pages 1–11, 2023.

Eugene F Douglass, Robert J Allaway, Bence Szalai, Wenyu Wang, Tingzhong Tian, Adrià Fernández-Torras, Ron Realubit, Charles Karan, Shuyu Zheng, Alberto Pessia, et al. A community challenge for a pancancer drug mechanism of action inference from perturbational profile data. Cell Reports Medicine, 3(1), 2022.

Mohamed Reda El Khili, Safyan Aman Memon, and Amin Emad. Marsy: a multitask deep-learning framework for prediction of drug combination synergy scores. Bioinformatics, 39(4):btad177, 2023.

Di He, Qiao Liu, You Wu, and Lei Xie. A context-aware deconfounding autoencoder for robust prediction of personalized clinical drug response from cell-line compound screening. Nature Machine Intelligence, 4(10):879–892, 2022.

Francesco Iorio, Theo A Knijnenburg, Daniel J Vis, Graham R Bignell, Michael P Menden, Michael Schubert, Nanne Aben, Emanuel Gonçalves, Syd Barthorpe, Howard Lightfoot, et al. A landscape of pharmacogenomic interactions in cancer. Cell, 166(3):740–754, 2016.

Joseph D Janizek, Safiye Celik, and Su-In Lee. Explainable machine learning prediction of synergistic drug combinations for precision cancer medicine. BioRxiv, page 331769, 2018.

Fredrik Johansson, Uri Shalit, and David Sontag. Learning representations for counterfactual inference. In International conference on machine learning, pages 3020–3029. PMLR, 2016.

Halil Ibrahim Kuru, Oznur Tastan, and A Ercument Cicek. Matchmaker: a deep learning framework for drug synergy prediction. IEEE/ACM transactions on computational biology and bioinformatics, 19(4):2334–2344, 2021.

Alexander Ling, Robert F Gruener, Jessica Fessler, and R Stephanie Huang. More than fishing for a cure: The promises and pitfalls of high throughput cancer cell line screens. Pharmacology & therapeutics, 191:178–189, 2018.

S Loewe. The problem of synergism and antagonism of combined drugs. Arzneimittel-forschung, 3(6):285–290, 1953.

Mohammad Lotfollahi, Anna Klimovskaia Susmelj, Carlo De Donno, Leon Hetzel, Yuge Ji, Ignacio L Ibarra, Sanjay R Srivatsan, Mohsen Naghipourfar, Riza M Daza, Beth Martin, et al. Predicting cellular responses to complex perturbations in high-throughput screens. Molecular Systems Biology, page e11517, 2023.

Thomas M Norman, Max A Horlbeck, Joseph M Replogle, Alex Y Ge, Albert Xu, Marco Jost, Luke A Gilbert, and Jonathan S Weissman. Exploring genetic interaction manifolds constructed from rich single-cell phenotypes. Science, 365(6455): 786–793, 2019.

Kristina Preuer, Richard PI Lewis, Sepp Hochreiter, Andreas Bender, Krishna C Bulusu, and Günter Klambauer. Deepsynergy: predicting anti-cancer drug synergy with deep learning. Bioinformatics, 34(9):1538–1546, 2018.

William C Reinhold, Margot Sunshine, Hongfang Liu, Sudhir Varma, Kurt W Kohn, Joel Morris, James Doroshow, and Yves Pommier. Cellminer: a web-based suite of genomic and pharmacologic tools to explore transcript and drug patterns in the nci-60 cell line set. Cancer research, 72(14):3499–3511, 2012.

Joseph M Replogle, Reuben A Saunders, Angela N Pogson, Jeffrey A Hussmann, Alexander Lenail, Alina Guna, Lauren Mascibroda, Eric J Wagner, Karen Adelman, Gila Lithwick-Yanai, et al. Mapping information-rich genotype-phenotype landscapes with genome-scale perturb-seq. Cell, 185(14):2559–2575, 2022.

Yusuf Roohani, Kexin Huang, and Jure Leskovec. Predicting transcriptional outcomes of novel multigene perturbations with gears. Nature Biotechnology, pages 1–9, 2023.

Donald B Rubin. Estimating causal effects of treatments in randomized and non-randomized studies. Journal of educational Psychology, 66(5):688, 1974.

Stefan Schrod, Andreas Schäfer, Stefan Solbrig, Robert Lohmayer, Wolfram Gronwald, Peter J Oefner, Tim Beißbarth, Rainer Spang, Helena U Zacharias, and Michael Altenbuchinger. Bites: balanced individual treatment effect for survival data. Bioinformatics, 38(Supplement 1):i60–i67, 2022.

Stefan Schrod, Fabian Sinz, and Michael Altenbuchinger. Adversarial distribution balancing for counterfactual reasoning. arXiv preprint arXiv:2311.16616, 2023.

Uri Shalit, Fredrik D Johansson, and David Sontag. Estimating individual treatment effect: generalization bounds and algorithms. In International conference on machine learning, pages 3076–3085. PMLR, 2017.

Hossein Sharifi-Noghabi, Parsa Alamzadeh Harjandi, Olga Zolotareva, Colin C Collins, and Martin Ester. Out-of-distribution generalization from labelled and unlabelled gene expression data for drug response prediction. Nature Machine Intelligence, 3(11):962–972, 2021.

Sanjay R Srivatsan, José L McFaline-Figueroa, Vijay Ramani, Lauren Saunders, Junyue Cao, Jonathan Packer, Hannah A Pliner, Dana L Jackson, Riza M Daza, Lena Christiansen, et al. Massively multiplex chemical transcriptomics at single-cell resolution. Science, 367(6473):45–51, 2020.

Aravind Subramanian, Rajiv Narayan, Steven M Corsello, David D Peck, Ted E Natoli, Xiaodong Lu, Joshua Gould, John F Davis, Andrew A Tubelli, Jacob K Asiedu, et al. A next generation connectivity map: L1000 platform and the first 1,000,000 profiles. Cell, 171(6):1437–1452, 2017.

Bhagwan Yadav, Krister Wennerberg, Tero Aittokallio, and Jing Tang. Searching for drug synergy in complex dose–response landscapes using an interaction potency model. Computational and structural biotechnology journal, 13:504–513, 2015.

Liuyi Yao, Sheng Li, Yaliang Li, Mengdi Huai, Jing Gao, and Aidong Zhang. Representation learning for treatment effect estimation from observational data. Advances in neural information processing systems, 31, 2018.

Chaoyang Ye, Daniel J Ho, Marilisa Neri, Chian Yang, Tripti Kulkarni, Ranjit Randhawa, Martin Henault, Nadezda Mostacci, Pierre Farmer, Steffen Renner, et al. Drug-seq for miniaturized high-throughput transcriptome profiling in drug discovery. Nature communications, 9(1):4307, 2018.

Jinsung Yoon, James Jordon, and Mihaela Van Der Schaar. Ganite: Estimation of individualized treatment effects using generative adversarial nets. In International conference on learning representations, 2018.

Bulat Zagidullin, Jehad Aldahdooh, Shuyu Zheng, Wenyu Wang, Yinyin Wang, Joseph Saad, Alina Malyutina, Mohieddin Jafari, Ziaurrehman Tanoli, Alberto Pessia, et al. Drugcomb: an integrative cancer drug combination data portal. Nucleic acids research, 47(W1):W43–W51, 2019.

