## Supplementary Materials for "CODEX: COunterfactual Deep learning for the *in-silico* EXploration of cancer cell line perturbations"

#### S1 Implementation details

CODEX is implemented using the PyTorch machine learning library. It uses a dense feed-forward architecture in combination with ReLU activation functions and Dropout layers. Further, it uses early-stopping to prevent overfitting and the minimal/maximal validation score is used to select the best performing model. Our implementation of CODEX is publicly available at `https://github.com/sschrod/CODEX`. All experiments were run on a cluster with 8 Nvidia A100 GPUs, 256 CPU cores and 512GB of RAM. Nevertheless, running the reconstruction experiments is computationally feasible for almost any modern GPU with CODEX requiring less than 5GB of GPU memory. Note that the drug-synergy experiments require more GPU memory due to the large treatment specific layers.

Table S1: CODEX hyper-parameter ranges for the drug-synergy experiment

| Parameter | Range |
|---|---|
| Dim. layers | $\{[4096, 2048, 1024, 512], [2048, 1024, 512, 256]\}$ |
| Learning Rate $\{0.0001\}$ Batch Size | $\{4096, 1024\}$ |
| Dropout | $\{0.2, 0.4\}$ |
| Weight Decay | $\{0.001, 0.01, 0.1, 1\}$ |

#### S2 The drug-synergy experiment

The hyper-parameter ranges used for the drug-synergy experiment are listed in Table S1. We performed an extensive grid-search, covering all possible combinations of the listed parameters. Following El Khili et al. [2023] and Preuer et al. [2018], we used a large architecture with many of nodes. The layers are split into two encoder layers, one treatment specific layer and one decoder layer, according to the "Dim. layers" parameters outlined in Table S1. The decoder is further connected to a single outcome node, representing the synergy score.

All models were trained until the validation MSE (10% of the training data) did not increase for 50 consecutive epochs. Finally, the best model was selected based on minimal validation MSE.

In addition to the results for the ZIP synergy score reported in the main manuscript, Table S2 shows the performance obtained for the $S_{\text{mean}}$ synergy score (following El Khili et al. [2023]).

Table S2: Mean performance of $S_{\text{mean}}$ synergy prediction on stratified 5-fold cross validation holding out specific drug pairs for testing.

| Model | Setting 1 | | | Setting2 | | |
|---|---|---|---|---|---|---|
| | SCC | PCC | RMSE | SCC | PCC | RMSE |
| CODEX | 0.820 (±.003) | 0.859 (±.003) | 8.59 (±.10) | 0.800 (±.005) | 0.838 (±.008) | 9.13 (±.37) |
| MARSY | 0.836 (±.002) | 0.864 (±.003) | 8.42 (±.09) | 0.809 (±.007) | 0.841 (±.011) | 9.06 (±.45) |
| MatchMaker | 0.810 (±.003) | 0.840 (±.005) | 9.84 (±.23) | 0.788 (±.009) | 0.816 (±.015) | 10.34 (±.46) |
| TreeCombo (XGBoost) | 0.817 (±.002) | 0.852 (±.002) | 8.77 (±.06) | 0.775 (±.010) | 0.815 (±.011) | 9.69 (±.37) |
| DeepSynergy | 0.803 (±.003) | 0.843 (±.002) | 9.14 (±.12) | 0.762 (±.004) | 0.804 (±.011) | 10.17 (±.42) |

Table S3: CODEX hyper-parameter ranges for the drug-dose and Combosciplex experiment

| Parameter | Range |
|---|---|
| Dim. layers | $\{[1024, 512, 256], [512, 256, 128]\}$ |
| Learning Rate $\{0.001, 0.0001\}$ Batch Size | $\{1024, 2048\}$ |
| Dropout | $\{0.0, 0.2\}$ |
| Weight Decay | $\{0.001, 0.00001, 0.0000001\}$ |

### S3 Drug-dose and Combosciplex experiments

Both, the drug-dose and Combosciplex experiments use the same set of hyper-parameters outlined in Table S3. Here, the dimension of the layers describes the 3 layer encoder, the latent representations consist of a single layer. The size of the last decoder layer and the decoder is given by the same 3 layer structure as the encoder, but in reversed order. For all reconstruction experiments, the decoder maps to a final outcome consisting of the mean and the variance of the post-perturbation data (the outcome has twice the size of the input features). Accordingly, the model was optimized using the Gaussian Negative-log-likelihood. Similar to the previous experiment, we performed a grid-search covering all possible parameter combinations and the best models were selected based on maximal reconstruction performance in terms of the validation coefficient of determination $R^2$ on the top 50 DEGs. For both experiments, we used 25% of the training data for validation and a patience of 50 epochs.

In addition to the individual dose-response curves, we report the median reconstruction performance for all doses (on validation and ood data) in Figure S1a and for the unseen dose (ood) in Figure S1b.

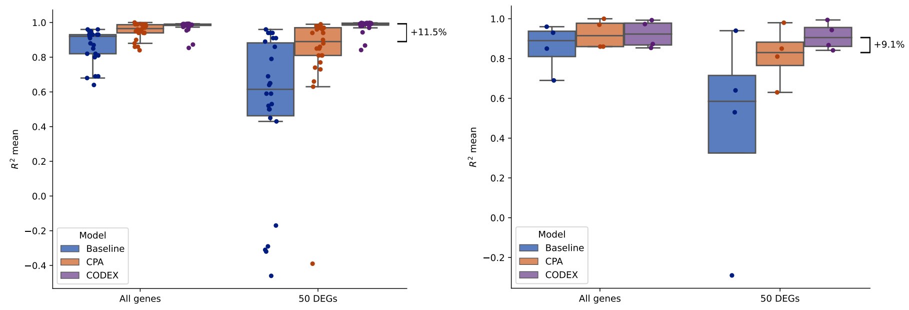


(a) Ood dosages. (b) Test and ood dosages.

Figure S1: Box plot for the reconstruction performance on Sciplex2 data.

Further, Table S4 shows all treatment combinations present in the Combosciplex data, with control samples highlighted in blue and ood combinations in red. Figure S2 shows the median reconstruction performance for the left out combinations.

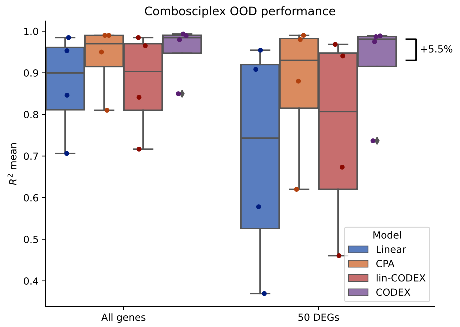


Figure S2: Box plot of the reconstruction performance of the four unobserved treatment combinations on Combosciplex data.

### S4 Gene-perturbation experiments

The experimental design of perturbation experiment on the Norman et al. [2019] follows the design of the drug-dose and Combosciplex experiments. The used hyper-parameters are listed in Table S5. In contrast to previous experiments, we evaluated the perturbation experiments with respect to PCCs and reconstruction MSE (following Roohani et al. [2023]). Hence, we selected the best performing model based on the minimal validation MSE for the top 20 DEGs.

In addition to the main experiments on Norman et al. [2019] data, we repeated the experiment of Lotfollahi et al. [2023]. Accordingly, we used 13

Table S4: List of all Combosciplex perturbation combinations, ood combinations are marked in red and control samples in blue

| Compound 1 | Compound 2 | Number of Cells |
|---|---|---|
| Dacinostat | PCI-34051 | 3302 |
| SRT3025 | Cediranib | 3017 |
| Givinostat | Cediranib | 2785 |
| DMSO | SRT2104 | 2757 |
| Givinostat | Curcumin | 2739 |
| Givinostat | Sorafenib | 2736 |
| Givinostat | Carmofur | 2694 |
| Givinostat | crizotinib | 2666 |
| Givinostat | dasatinib | 2421 |
| Givinostat | SRT2104 | 2357 |
| DMSO | dasatinib | 2343 |
| Givinostat | SRT1720 | 2265 |
| panobinostat | Curcumin | 2244 |
| Cediranib | PCI-34501 | 2166 |
| panobinostat | Sorafenib | 2013 |
| panobinostat | SRT2104 | 1974 |
| panobinostat | dasatinib | 1956 |
| Dacinostat | Danusertib | 1941 |
| panobinostat | SRT3025 | 1890 |
| DMSO | Dacinostat | 1869 |
| panobinostat | SRT1720 | 1828 |
| panobinostat | PCI-34051 | 1814 |
| DMSO | Givinostat | 1685 |
| panobinostat | crizotinib | 1643 |
| DMSO | panobinostat | 1578 |
| DMSO | DMSO | 1451 |
| Givinostat | Tanespimycin | 1314 |
| Dacinostat | dasatinib | 1231 |
| panobinostat | Alvespimycin | 996 |
| DMSO | Alvespimycin | 758 |
| SRT2104 | Alvespimycin | 520 |
| Alvespimycin | Pirarubicin | 477 |
| | | 63430 |

Table S5: CODEX hyper-parameter ranges for the perturbation experiments

| Parameter | Range |
|---|---|
| Dim. layers | $\{[1024, 512, 256], [512, 128, 64]\}$ |
| Learning Rate $\{0.001, 0.0001\}$ Batch Size | $\{1024, 512, 256\}$ |
| Dropout | $\{0.1, 0.2, 0.4\}$ |
| Weight Decay | $\{0.00001, 0.000001, 0.0000001\}$ |

train/test splits with $\sim$ 10 perturbation combinations left out in each split, covering all of the 131 two gene perturbations. Figure S3 shows the $R^2$ reconstruction performance for all genes and the top 50 DEGs, comparing CODEX to CPA Lotfollahi et al. [2023], to the linear baseline and to the random baseline.

Following the results on the Norman et al. [2019] data, we evaluated the Replogle et al. [2022] data for the set of parameters that was most frequently selected. Hence, we used the architecture with a decoder consisting of $[512, 128, 64]$ nodes, a learning rate of 0.001, a batch site of 256, a dropout rate of 0.1, and weight decay of $10^{-6}$.

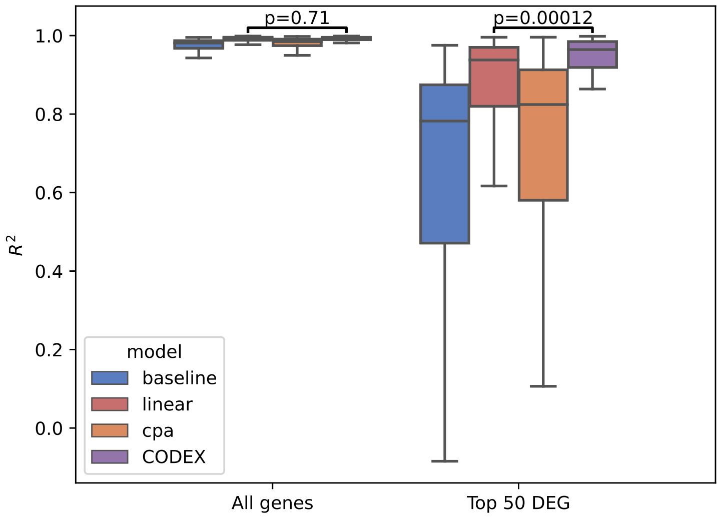


Figure S3: Reconstruction performance in terms of $R^2$ for all genes (left) and the top 50 DEGs (right) in the experimental setting of Lotfollahi et al. [2023].

### References


Mohamed Reda El Khili, Safyan Aman Memon, and Amin Emad. Marsy: a multitask deep-learning framework for prediction of drug combination synergy scores. *Bioinformatics*, 39(4):btad177, 2023.

Mohammad Lotfollahi, Anna Klimovskaia Susmelj, Carlo De Donno, Leon Hetzel, Yuge Ji, Ignacio L Ibarra, Sanjay R Srivatsan, Mohsen Naghipourfar, Riza M Daza, Beth Martin, et al. Predicting cellular responses to complex perturbations in high-throughput screens. *Molecular Systems Biology*, page e11517, 2023.

Thomas M Norman, Max A Horlbeck, Joseph M Replogle, Alex Y Ge, Albert Xu, Marco Jost, Luke A Gilbert, and Jonathan S Weissman. Exploring genetic interaction manifolds constructed from rich single-cell phenotypes. *Science*, 365(6455):786–793, 2019.

Kristina Preuer, Richard PI Lewis, Sepp Hochreiter, Andreas Bender, Krishna C Bulusu, and Günter Klambauer. Deepsynergy: predicting anti-cancer drug synergy with deep learning. *Bioinformatics*, 34(9):1538–1546, 2018.

Joseph M Replogle, Reuben A Saunders, Angela N Pogson, Jeffrey A Hussmann, Alexander Lenail, Alina Guna, Lauren Mascibroda, Eric J Wagner, Karen

Adelman, Gila Lithwick-Yanai, et al. Mapping information-rich genotype-phenotype landscapes with genome-scale perturb-seq. *Cell*, 185(14):2559–2575, 2022.

Yusuf Roohani, Kexin Huang, and Jure Leskovec. Predicting transcriptional outcomes of novel multigene perturbations with gears. *Nature Biotechnology*, pages 1–9, 2023.
